# Dynamics of BMP signaling in the early *Drosophila* embryo

**DOI:** 10.1101/2022.10.20.513072

**Authors:** Hadel Y. Al Asafen, Aydin Beseli, Sharva Hiremath, Cranos M. Williams, Gregory T. Reeves

## Abstract

In developing tissues, morphogen gradients are thought to initialize gene expression patterns. However, the relationship between the dynamics of morphogen-encoded signals and gene expression decisions are largely unknown. Here we examine the dynamics of the Bone Morphogenetic Protein (BMP) pathway in *Drosophila* blastoderm-stage embryos. In this tissue, the BMP pathway is highly dynamic: it begins as a broad and weak signal on the dorsal half of the embryo, then 20-30 min later refines into a narrow, intense peak centered on the dorsal midline. This dynamical progression of the BMP signal raises questions of how it stably activates target genes. Therefore, we performed live imaging of the BMP signal and found that dorsal-lateral cells experience only a short transient in BMP signaling, after which the signal is lost completely. Moreover, we measured the transcriptional response of the BMP target gene *pannier* in live embryos and found it to remain activated in dorsal-lateral cells, even after the BMP signal is lost. Our findings may suggest that the BMP pathway activates a memory, or “ratchet” mechanism that may sustain gene expression.

## Introduction

In developing tissues, gene expression decisions are initiated by morphogen gradients, which are often assumed to deliver a continuous input of a steady signal (Ashe and Briscoe, 2006; Dresch et al., 2013; Gurdon and Bourillot, 2001; Jaeger et al., 2004; Reeves et al., 2005; Wharton et al., 2004). However, the dynamics of morphogen gradients may also play a role in gene expression (Delotto et al., 2007; Gregor et al., 2007; Nahmad and Stathopoulos, 2009; Reeves et al., 2012; Ribes et al., 2010). In the past two decades, studies using GFP-tagged morphogens – including early *Drosophila* morphogens Bicoid and Dorsal; Dpp in the wing imaginal disc; and the Hedgehog, Wnt, and TGF-β families in vertebrates – have revealed that the establishment of morphogen gradients is a highly dynamic and complex process (Delotto et al., 2007; Entchev et al., 2000; Gregor et al., 2007; Holzer et al., 2012; Luz et al., 2014; Reeves et al., 2012; Ribes et al., 2010; Teleman and Cohen, 2000; Wallkamm et al., 2014; Wartlick et al., 2011; Williams et al., 2004; Zhou et al., 2012). Live imaging studies – of both morphogen input and the resulting gene expression output – have challenged the established view that tissue patterning relies on a constant, steady-state level of signaling to regulate gene expression (Aldaz et al., 2010; Ashe and Briscoe, 2006; Dresch et al., 2013; Gurdon and Bourillot, 2001; Gurdon et al., 1995; Jaeger et al., 2004; Mavrakis et al., 2010; Oates et al., 2009; Pantazis and Supatto, 2014; Reeves et al., 2005; Wharton et al., 2004).

The Bone Morphogenetic Protein (BMP) signaling pathway in the blastoderm stage *Drosophila* embryo is a classic example of a highly dynamic morphogen-mediated system. BMPs are a group of signaling molecules that were initially discovered for their ability to induce bone formation. BMPs are now known to play crucial roles in many tissue patterning processes, including establishing the dorsal-most tissues during *Drosophila* embryogenesis. Roughly two hours after fertilization, the nuclei have completed 13 nuclear division cycles, and roughly 6000 nuclei are present at the embryo’s cortex. During the 14^th^ interphase [nuclear cycle (nc) 14], BMP signaling acts through two of the *Drosophila* BMP ligands, Dpp (expressed on the dorsal 50% of the embryo) and Screw (Scw; uniformly expressed) (Arora et al., 1994; Padgett et al., 1987). The ligand-bound Type I receptors, Thickveins and Saxophone (Brummel et al., 1994), together with Punt (a Type II receptor) (Letsou et al., 1995), phosphorylate Mothers against Dpp (Mad; homolog of vertebrate Smad1), which enters the nucleus with Co-Smad Medea (Med; homolog of Smad4) to regulate gene expression (Fig. 1A) (Raftery and Sutherland, 2003; Raftery et al., 1995).

**Figure 1.**
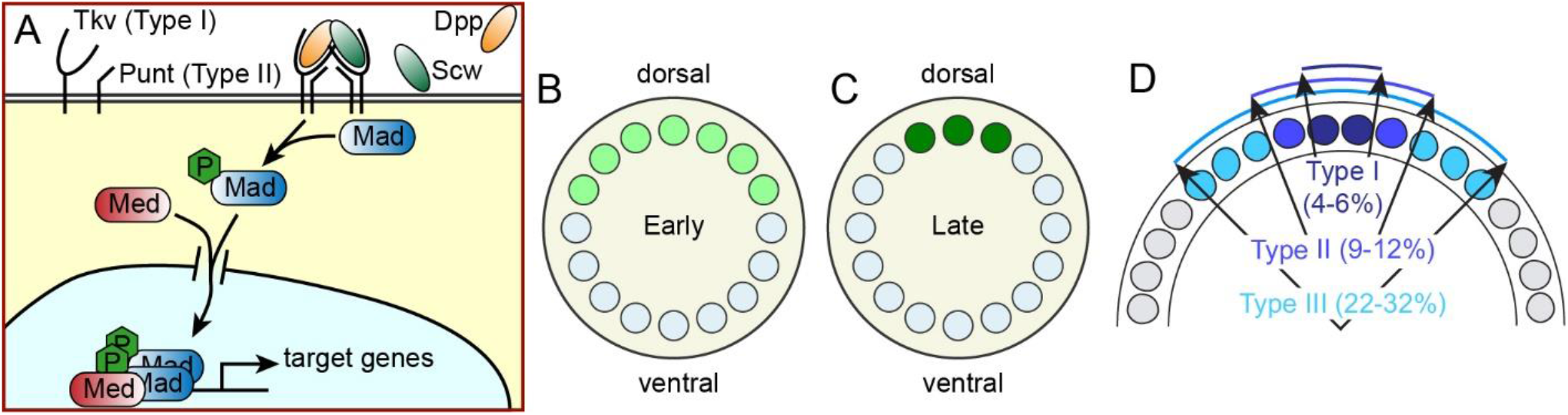
Schematic of the BMP network. **(A)** BMP ligands Dpp and Scw bind to their cognate Type I receptors, which dimerize with Type II receptors. The active receptor signaling complex causes the phosphorylation of Mad, which partners with Med to enter the nucleus. **(B)** The early stage of BMP signaling is broad and weak. **(C)** Roughly 20 min later, BMP signaling has intensified into a narrow domain. **(D)** BMP target genes, denoted Types I-III, are expressed in three nested domains. The ventral boundaries of these domains are noted on the figure.

The spatial pattern of BMP signaling is highly dynamic, and has been assayed by immunostaining of the phosphorylated form of Mad (pMad) in fixed embryos (Dorfman and Shilo, 2001; Gavin-Smyth et al., 2013; O’Connor et al., 2006; Raftery and Sutherland, 2003; Ross et al., 2001; Rushlow, 2001; Tanimoto et al., 2000; Wang and Ferguson, 2005). In early nc 14, the initial pattern of BMP signaling is broad and weak, covering the dorsal 30-40% of the embryo (Fig. 1B). Just 20-30 minutes later, the BMP signal intensifies and sharpens to a narrow domain in the dorsal-most 5-6% of the embryo (Fig. 1C) (Gavin-Smyth et al., 2013; O’Connor et al., 2006; Raftery and Sutherland, 2003). The broad, low intensity signal likely directs the dorsal half of the embryo to become dorsal epidermis, and the cells in the domain of late, high intensity signal eventually develop into the amnioserosa (Bier and De Robertis, 2015).

How cells interpret this dynamic BMP signal to form stable gene expression patterns, and whether the precise progression of BMP dynamics is necessary for proper development, is unknown. Three nested domains of BMP target genes (Types I-III) have been observed (Fig. 1D) (Ashe et al., 2000; Liang et al., 2012; Lin et al., 2006; Xu et al., 2005). However, at no single stage do quantitative data suggest the BMP profile has the ability to simultaneously specify all three domains: the early, broad domain is too weak to activate Type I genes, while the refined state is too narrow to maintain Type III genes (Gavin-Smyth et al., 2013). Moreover, detailed studies of the regulatory units of BMP target genes have raised additional questions, as no clear affinity-based enhancer structure has been found (Liang et al., 2012; Lin et al., 2006; Xu et al., 2005). In particular, qualitative studies have noted that the Type III domain is not clearly correlated with the of pMad domain (Xu et al., 2005).

The best-studied Type III gene, *pannier* (*pnr*), has a complex expression pattern restricted to the 25% dorsal-most cells that lie between 40% and 80% AP (0% AP being the anterior pole) (Liang et al., 2012). *pnr* expression is composed of two distinctly-regulated domains: a contiguous “dorsal patch” of cells (Liang et al., 2012), and six AP-modulated stripes overlaid on the patch. Regulation of the patch region is partially separable from the stripes, as an enhancer for *pnr* was isolated from the first intron of *pnr*. This enhancer, denoted the *pnr P3* enhancer, drives the expression of the dorsal patch and two of the six stripes (Liang et al., 2012). The patch requires BMP signaling, as it is lost in *dpp* mutants. The stripes become dramatically weaker in these mutants as well (Liang et al., 2012).

In this paper, we measured the dynamics of both BMP signaling and *pnr* expression in live embryos. We first showed in fixed embryos that nuclear localization of Med-GFP is an appropriate assay for BMP signaling, as it correlates strongly with pMad staining. Next, we imaged the dynamics of Med-GFP in live embryos and showed that cells in the tails of the Type III domain (between 9% and 25% DV coordinate) experience only a transient of BMP signaling in the first few minutes of nc 14. Finally, we showed in both live and fixed embryos that the cells in this same domain continue to express *pnr* throughout nc 14. We argue that *pnr* requires BMP for activation, but not maintenance. We hypothesize this may constitute a ratchet module, similar to that found in Hedgehog signaling in the wing disc or Dorsal activity in DV patterning of the early embryo (Irizarry et al., 2020; Nahmad and Stathopoulos, 2009).

## Results and Discussion

### Nuclear Med-GFP mirrors pMad

In cells with activated BMP signaling, Mad is phosphorylated and both pMad and Med enter the nucleus (Fig. 1A). To determine whether nuclear localization of Med-GFP is an appropriate assay for BMP signaling in live embryos, we first measured pMad and nuclear Med-GFP in fixed, nc 14 embryos (see Methods). Nuclear localized Med-GFP was higher on the dorsal side, in the same nuclei as the pMad staining (Fig. 2A,B). While nuclear Med-GFP was not necessarily higher than cytoplasmic Med-GFP in dorsal nuclei, in the rest of the embryo Med-GFP was excluded from the nuclei. Late Stage 5/early Stage 6 embryos do in fact exhibit higher levels of nuclear Med-GFP than the surrounding cytoplasm (Sutherland et al., 2003) (Fig. S1).

**Figure 2.**
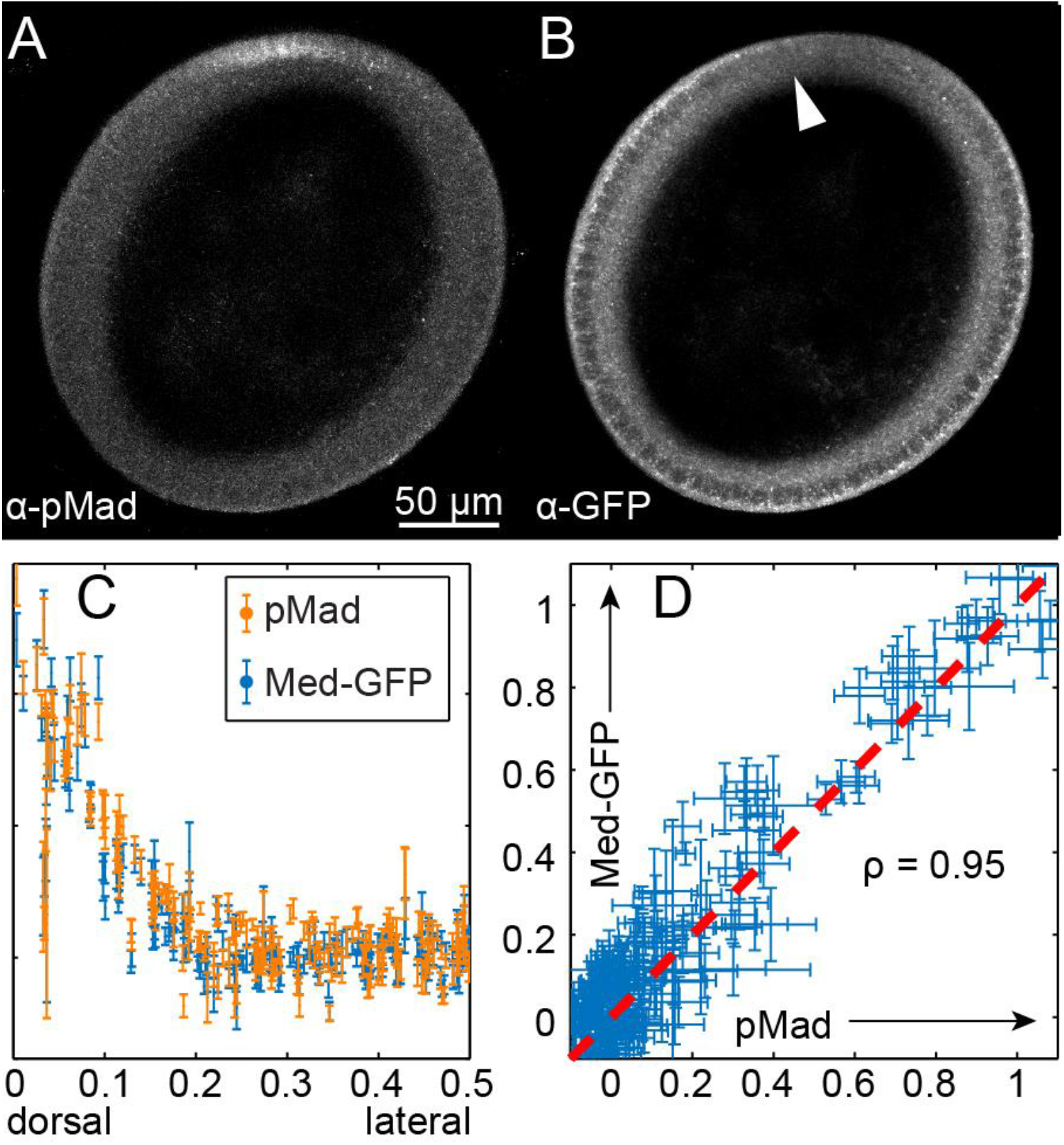
Correlation between pMad and nuclear Med-GFP. **(A,B)** Cross section of nc 14 embryo expressing Med-GFP and immunostained against pMad (A) and GFP (B). **(C)** Quantification of nuclear intensities in embryo shown in (A,B). The nuclear intensities of pMad and Med-GFP are highly correlated. **(D)** Plot of the Med-GFP nuclear intensity versus the pMad nuclear intensity. Errorbars indicate S.E.M.

Our quantitative analysis showed a strong overlap between pMad and the nuclear intensity of Med-GFP (Figs. 2C, S2). The high correlation (ρ = 0.9± 0.05 s.d.) becomes clear when the normalized intensity of Med-GFP is plotted against that of pMad for each nucleus (Figs. 2D, S2; see Supplemental Methods for normalization procedure). Therefore, these data in fixed embryos support using nuclear localization of Med-GFP in live embryos as a readout of BMP signaling.

### Measurements of nuclear Med-GFP dynamics

To measure the dynamics of nuclear Med-GFP, we imaged live embryos (n = 2) end-on with a confocal microscope roughly 100 μm from the posterior pole (Figs. 3A-C) (Carrell and Reeves, 2015). Expression of H2A-RFP allowed us to segment the nuclei to distinguish nuclear from cytoplasmic pixels. The Med-GFP gradient was initially broad and weak, then refined into a narrow, intense peak before gastrulation (Figs. 3D-F; Movies S1,2). To model the nature of the BMP transient, especially at the location of the Type III boundary (∼25% DV), we averaged data from the two embryos (see Supplemental Methods and Movie S3). These quantitative measurements captured crucial aspects the BMP signaling dynamics that were previously unknown, including the single-embryo dynamics of gradient amplitude and width, as well as tracking of the transient at specific DV coordinates. In particular, we found the early peak was wide enough to activate the Type III domain, yet the gradient at later time points was not (Fig. 3G). In its most refined state, the gradient was wide enough to plausibly activate genes in both the Type I and II domains.

**Figure 3:**
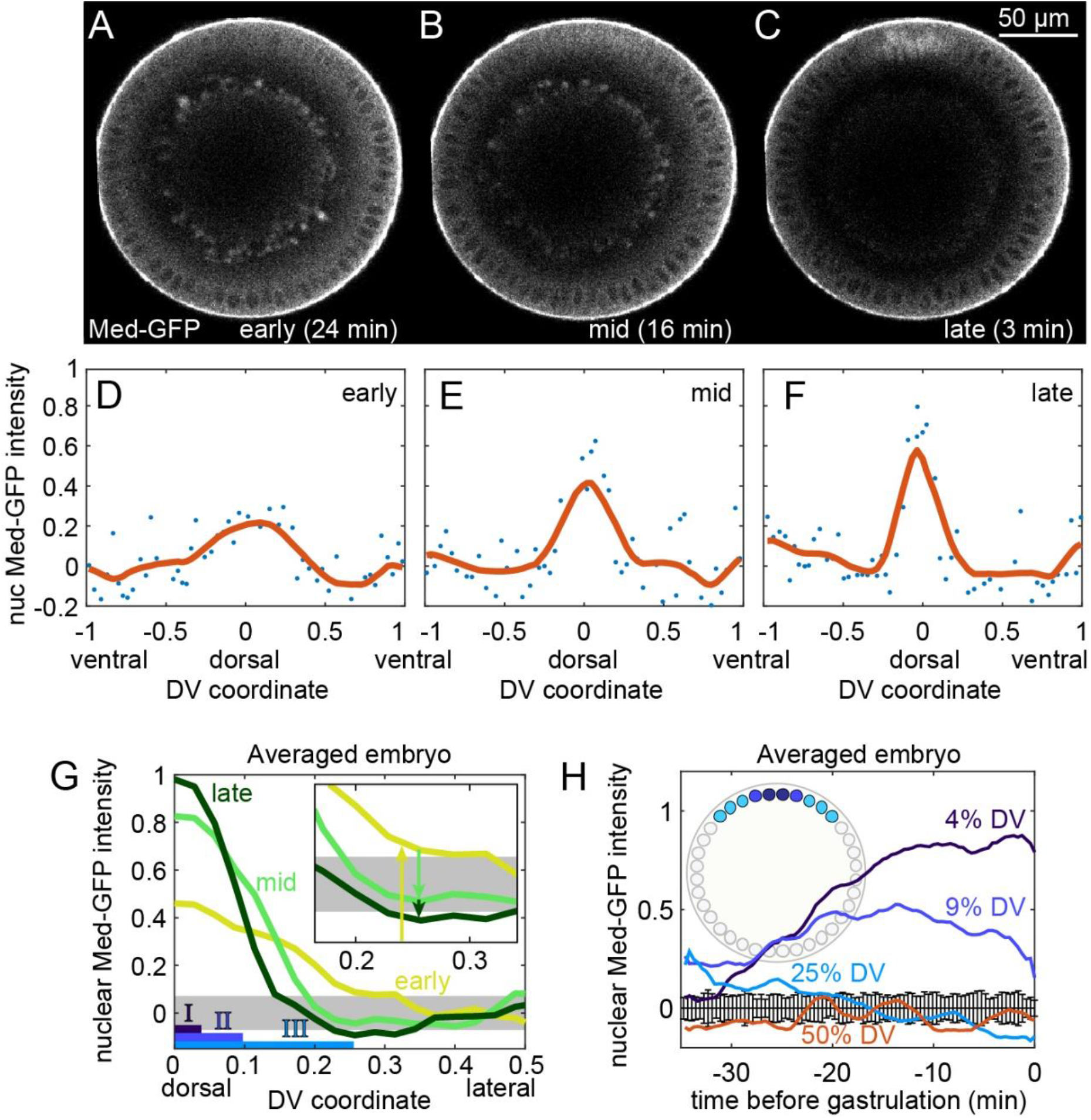
Dynamics of Med-GFP. **(A-C)** Snapshots of live embryo expressing Med-GFP. Time stamps for early, mid, late: time before gastrulation. **(D-F)** Quantification of the nuclear Med-GFP gradient from (A-C). **(G)** Plots of early, mid, and late gradients from an averaged embryo. The extents of BMP target gene Types I-III are illustrated at the bottom. Gray rectangle represents uncertainties of background levels derived from control embryos that express H2A-RFP but not Med-GFP. **(H)** Time course of BMP signaling for different locations along the DV axis. Errorbars centered around zero represent the S.E.M. of control embryos that express H2A-RFP but not Med-GFP.

Nuclei that lay at the boundaries of Type I-III domains experienced distinct BMP dynamics. Nuclear Med-GFP intensities at 4% DV (Type I boundary) rose continuously (Fig. 3H). In contrast, nuclei between the Type II and III boundaries experienced a transient in GFP intensity. At 9% DV (Type II boundary), GFP intensity peaked roughly 15 mins before gastrulation, followed by a gradual decrease as the gradient continued to refine. The transient intensity at the Type III boundary (25% DV) was even shorter: nuclear Med-GFP levels were above the background levels only briefly, then dropped to a level indistinguishable from background levels roughly 25 min before gastrulation. Background levels were determined by control embryos expressing H2A-RFP but not Med-GFP (see Supplemental Methods and Fig. S3). Thus, cells beyond the Type II domain boundary experienced BMP signaling transiently, which raised the question of the fate of Type III transcription in cells between 9% and 25% DV.

### pnr is transcribed in cells that lack BMP signaling

To measure the extent and timing of Type III gene expression, we first detected nascent transcripts of *pnr* in fixed embryos using RNA probes that hybridize to the first intron of *pnr* (Fig. 4A). We co-immunostained these embryos with an antibody against pMad. In Stage 6 embryos, when the cephalic furrow was evident, the pMad stripe had become narrow and intense, while the domain of active *pnr* expression remained wide. These results suggested that a short transient of BMP signaling is sufficient to activate long-term *pnr* expression. However, the *pnr* domain is complex, comprising at least two overlapping sub-patterns: the dorsal patch and a six-striped pattern (Liang et al., 2012). While the ventral boundary of the six-striped pattern is set by the activity of the repressor Brinker (Brk) (Jaźwińska et al., 1999a, 1999b), which is expressed in a broad lateral stripe between ∼50% and 80% DV (Fig. S4), BMP signaling sets the boundary for the dorsal patch.

**Figure 4:**
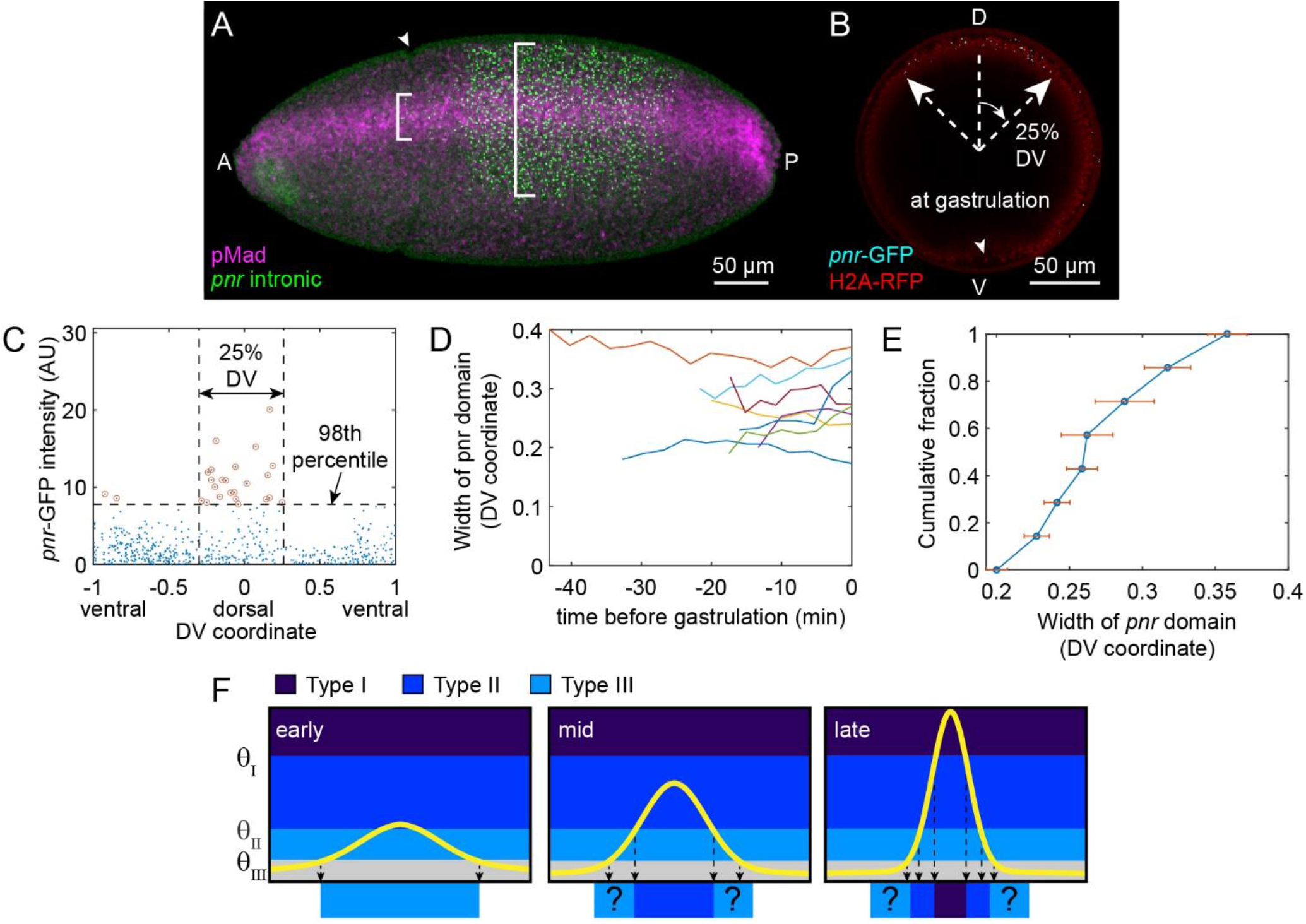
Transcription of Type III gene *pnr*. **(A)** Fixed embryo immunostained against pMad (magenta) and hybridized with an antisense probe against the first intron of *pnr* (green). At this stage, the pMad signal has refined, but the *pnr* extent remains wide (note white brackets). The cephalic furrow can be seen (arrowhead), indicating the beginning of gastrulation. **(B)** Snapshot of live embryo expressing the *pnr-*GFP constructs (cyan dots), as well as H2A-RFP (red). The domain of cells actively transcribing from the *pnr P3* enhancer is 25% DV. The ventral furrow can be seen at this time point (arrowhead), indicating the beginning of gastrulation. **(C)** Plot of the *pnr-*GFP dot intensities at the time point shown in (B). Dots are counted as *pnr P3* transcription if they exceed the 98^th^ percentile. A 25% DV window captures most of these dots (see Supplementary Methods). **(D)** Quantification of the width of the *pnr-*GFP domain from eight embryo time courses. **(E)** Cumulative distribution plot of *pnr-*GFP domain widths. Points and errorbars are averages and S.E.M., respectively, across each time course. **(F)** Plausible model of the relationship between BMP signaling dynamics and gene expression. Early BMP signaling in above only the threshold, θ_III_, for Type III genes. At an intermediate stage, the signal intensifies to also activate Type II genes. At late stages, the signal becomes intense enough to activate both Types I and II. However, the location where the signal surpasses the Type III threshold is much narrower than the Type III domain (see vertical down arrows). The question marks represent a putative memory mechanism.

To specifically measure the dynamics of expression in the BMP-dependent dorsal patch, we used the MS2 system in live embryos (Bertrand et al., 1998; Forrest and Gavis, 2003; Forrest et al., 2004; Garcia et al., 2013; Lucas et al., 2013; Weil et al., 2006). We chose to drive expression of a reporter gene, *lacZ*, by the *pnr P3* enhancer fragment, which directs expression primarily of the dorsal patch, but also includes two of the six stripes (Liang et al., 2012). In BMP mutants, the dorsal patch driven by the *P3* enhancer is lost (Liang et al., 2012). Therefore, we chose to analyze the *pnr P3* enhancer because of its specificity for BMP signaling. We placed 24x MS2 sequences in the 5’UTR of the reporter construct. For convenience, we refer to the *pnr P3-MS2-lacZ*/MCP-GFP combination as *pnr-*GFP. We imaged these live embryos (also expressing H2A-RFP) end-on, which revealed nuclear dots of nascent transcripts (transcription in progress) of *pnr*-GFP in a wide domain consistent with the Type III domain, even at gastrulation (Fig. 4B). We quantified the width of the expression domain by determining the fraction of the DV axis that contains the highest number nuclear dots above the 98^th^ percentile of intensity while leaving out those below (Fig. 4C; see Supplementary Methods for more details). After quantifying the *pnr-*GFP expression domain in multiple embryos (n = 8; see Fig. 4D-E and Supplementary Movies 4-11), we found that the domain width does not systematically change over the course of nc 14 (Fig. 4D). These data show that the *pnr* dorsal patch maintains active gene expression in the Type III broad domain (27% ± 5% DV; Fig. 4E) until gastrulation begins. The transient nature of BMP signaling in these cells, together with the continued expression of *pnr*, could plausibly imply that some Type III genes require BMP signaling for activation, but not maintenance, which may constitute a memory, or “ratchet” mechanism (Fig. 4F).

It has been shown that other factors regulate *pnr* expression (Liang et al., 2012). In particular, Brk at least partially sets the boundary of *pnr*; however, there is also a portion of *pnr* expression that (1) requires BMP for activation (Winick et al., 1993), and (2) is not significantly regulated by endogenously-expressed Brk. Our results suggest that, at a minimum, the dorsal patch, largely driven by the *pnr P3* enhancer, is regulated by a memory mechanism.

This mechanism, in which the precise BMP dynamics are responsible for the proper pattern, could be similar to Hedgehog (Hh) signaling in the *Drosophila* wing disc. In that system, Hh signaling initially expands across ∼10 cell widths (Nahmad and Stathopoulos, 2009; Strigini and Cohen, 1997), but later refines due to a feedback mechanism. The cells that receive transient exposure of Hh signaling are able to permanently activate *dpp* expression. On the other hand, the receptor *patched* requires sustained Hh signaling, and is thus expressed in a narrow domain (Nahmad and Stathopoulos, 2009). Furthermore, a ratchet mechanism has recently been proposed for Dorsal-dependent activation of Twist on the ventral side of the embryos (Irizarry et al., 2020). Given these additional cases in which a memory mechanism may play a direct part in patterning, we propose this mechanism may be more widespread than previously appreciated.

## Methods

### Fly lines

The following fly lines were obtained from the Bloomington Drosophila Stock Center: His2AV-RFP (w[*]; P{w[+mC]=His2Av-mRFP1}II.2 -- BS# 23651), His2AV-RFP; MCP-GFP (y[1] w[*]; P{w[+mC]=His2Av-mRFP1}II.2; P{w[+mC]=nos-MCP.EGFP}2 -- BS# 60340), *nos*-Gal4 (w[1118]; P{w[+mC]=GAL4::VP16-nos.UTR}CG6325[MVD1] – BS# 4937). The UAS-Med-GFP line was a kind gift from Laurel Raftery (Miles et al., 2008).

### Creation of pnr P3-24x MS2-lacZ construct

The Pnr-P3 enhancer region was amplified from genomic DNA of wildtype flies. These sequences were cloned into the injection vector piB-hbP2-P2P-MS2-24x-lacZ-αTub3’UTR (a kind gift from Thomas Gregor)(Garcia et al., 2013) by replacing the hbP2 enhancer with the pnr p3 enhancer and the hb P2 promoter with an eve minimal promoter. These regulatory sequences in the plasmid drove the expression of lacZ, which contained 24 copies of the MS2 stem loops in its 5’UTR. This vector was injected into the 38F1 fly line (Genetivision) for recombination-mediated cassette exchange (RMCE)(Bateman et al., 2006).

### Fixation and immunostaining of embryos

Wildtype (laboratory strain yw) or nos-Gal4/UAS-Med-GFP embryos were aged to 2-4 hours after egg lay, then fixed in 37% formaldehyde according to published protocols (Kosman et al., 2004). Fluorescent in situ hybridization and fluorescent immunostaining were performed according to published protocols (Kosman et al., 2004).

Primary antibodies used were rabbit anti-Human Phospho-Smad2/3 (R&D Systems MAB8935, 1:500 dilution), goat anti-histone (Abcam ab12079, 1:100 dilution), goat anti-GFP (Rockland 600-101-215, 1:400 dilution), and goat anti-fluorescein (Rockland 600-101-096, 1:500 dilution; to detect the *pnr* intronic probe). In embryos co-stained for pMad and GFP, nuclei were detected with DAPI.

Secondary antibodies used were donkey anti-rabbit-Alexa Fluor 546 (Invitrogen A10040, 1:500 dilution) and donkey anti-goat conjugated with Alexa Fluor 647 (Invitrogen A21447, 1:500 dilution).

### Imaging fixed embryos

Nc 14 embryos expressing Med-GFP and immunostained against pMad and GFP were cross sectioned using a razor by removing the anterior and posterior thirds of the embryo, as described previously (Carrell and Reeves, 2015). The cross sections were mounted in 70% glycerol and imaged using a Zeiss LSM 710 microscope at 20x magnification. Z-stacks were taken with 15-25 z-slices at 2.5 μm intervals.

### Analysis of fixed embryos

The analysis of fixed embryo images followed our previously-published protocols (Trisnadi et al., 2013). Briefly, the border of the embryo was detected computationally, the nuclei were segmented, and for each nucleus, the mean intensity and standard error of the mean intensity of the pMad channel (Figs. 1,2) and Med-GFP channel (Fig. 2) were recorded. The position of each nucleus along the DV axis was also recorded. See Supplementary Methods for more information.

### Imaging live embryos

To image Med-GFP, *yw*; *His2AV-RFP*; *nos-Gal4* flies were crossed to *UAS-Med-GFP* flies. F1 females carrying all three constructs were then placed in embryo collection cages with males. The progeny embryos were collected for one hour on grape juice agar plates, dechorionated using bleach for 30 seconds, then rinsed using DI water. To prevent drying during mounting, the embryos were placed onto an agar plate. These embryos were then transferred to the short edge of a 22 mm coverslip, to which heptane glue (Supatto et al., 2009) had been applied, as described previously (Carrell and Reeves, 2015). Briefly, each embryo was oriented perpendicular to the short edge of the coverslip, such that approximately half of the embryo was adhered to the heptane glue on the coverslip and half was hanging off. The coverslip was then adhered with double-sided tape to a 21.5 mm-tall steel mounting block in the shape of a half cylinder with a 30 mm diameter (Carrell and Reeves, 2015). The short edge of the coverslip to which embryos were not adhered was aligned with the top of the mounting block, such that 0.5 mm of the coverslip (to which embryos were adhered) was extending past the bottom of the mounting block (Carrell and Reeves, 2015). The mounting block was then placed into a glass-bottom 35 mm petri dish with No. 1.5 coverslip (Mattek P35G-1.5-14-C), centered such that the 0.5 mm overhang of the coverslip was inside the cut bottom of the petri dish. DI water was added to the dish to prevent embryos from drying out. The glass-bottom dish/mounting block was placed on the Zeiss LSM 710 inverted microscope to collect images. A 40x 1.1 NA water immersion lens with 0.6 mm working distance was used. The long working distance allowed for imaging optical cross sections of the embryos > 100 μm away from the coverslip. The mounting process took roughly one hour so that embryos were at or near nc 14 upon the beginning of imaging.

To image *pnr*-GFP, virgin females of yw; His2AV-RFP; MCP-GFP flies were placed in embryo collection cages with males with *pnr-P3* enhancer 24xMS2-*lacZ* construct. The progeny embryos were subject to the same mounting and imaging procedure as the Med-GFP embryos, with the exception that 20 micron z-stacks (21 slices, 1 micron per slice) were taken at distances of 75-95 μm from the pole.

### Analysis of images of live Med-GFP embryos

Time course images of embryos expressing Med-GFP and H2A-RFP were analyzed in the following manner. First, in each frame (time point), the border of the embryo was detected. Next, the H2A-RFP image at each frame (time point) was segmented to identify the nuclei. To correct for intensity variations around the periphery of the embryo, the nuclear intensities or RFP, which are assumed to be uniform in the embryo, were used to create a normalization image. The nuclear intensities of Med-GFP were then calculated and normalized with the normalization image and with a laser calibration measurement (see Supplementary Methods for more information).

### Analysis of images of live pnr-GFP embryos

To detect spots of *pnr*-GFP in live embryos, we found the max intensity pixel in each nucleus in the GFP channel, subtracted it from the mean nuclear intensity, and normalized it by the intensity in the H2A-RFP channel.

The nuclei with the top 2% of values of *Y* at each time point clustered on the dorsal ∼30% of the embryo (Fig. 4) and were considered to be expressing *pnr*-GFP. For more information, including how the borders of the region of the embryo where *pnr*-GFP was expressed were determined, see Supplementary Methods.

